## Supplementary Data for "Autonomous computational prioritisation of colorectal cancer vulnerabilities via multi-scale AI swarms"

### Supplementary Information

Article Title: Autonomous computational prioritisation of colorectal cancer vulnerabilities via multi-scale AI swarms
Journal: Nature Computational Science
Authors: Christopher Baker et al.

#### Table of Contents


1. Supplementary Note 1: Algorithmic World Model Configuration (XGBoost)
2. Supplementary Note 2: Agentic Orchestration and LLM Prompting Strategy
3. Supplementary Note 3: In Silico Feature Ablation State-Space Perturbation Mathematics
4. Supplementary Table 1: Top 50 High-Variance Target Pool (CCLE Bowel Cohort)
5. Supplementary Note 4: Autonomous Agent Audit Trace (JSON Output)


#### Supplementary Note 1: Algorithmic World Model Configuration (XGBoost)

To map the in vitro baseline of 5-Fluorouracil resistance without overfitting to sample-specific noise, the digital twin predictive environment was constructed using XGBoost. The tree topology and regularisation parameters were strictly constrained to enforce the learning of broad biological pathway dynamics. The hyperparameters were defined programmatically in the pipeline as follows:

- Objective Function: reg:squarederror
- Number of Estimators (n_estimators): 100
- Learning Rate (learning_rate): 0.05
- Maximum Tree Depth (max_depth): 1
- Subsample Ratio (subsample): 0.8
- Column Sample by Tree (colsample_bytree): 0.8
- Minimum Child Weight (min_child_weight): 2 (Prevents isolated leaf nodes)
- L1 Regularisation (reg_alpha): 0.1
- L2 Regularisation (reg_lambda): 1.0

The model achieved an out-of-fold generalisation score of R^2 = 0.50 across the bowel lineage cohort prior to SHAP extraction.

#### Supplementary Note 2: Agentic Orchestration and LLM Prompting Strategy

To operate within strict clinical data governance requirements, the system utilised a quantised 4-bit Gemma-4 foundation model executed locally via llama.cpp. To bypass the inherent JSON parsing degradation and "reasoning loop" hallucinations commonly observed in quantised models, the CrewAI orchestrator was restricted using single-line conversational templates and exact stop-token matching (stop=["<end_of_turn>"]).

The system prompts did not rely on heuristic constraints but rather forced the Large Language Model (LLM) to interface strictly with the deterministic mathematical outputs.

##### Task 1: The Molecular Biologist Prompt

"Our multi-scale intersection pipeline evaluated [TARGET_GENE] for its role in 5-Fluorouracil response. DETERMINISTIC GUARDRAILS: 1. Human Survival Direction: [ACCELERATED MORTALITY / IMPROVED SURVIVAL] 2. CRITICAL GUARDRAIL: If the PDX data shows enhanced tumour shrinkage in the High expression cohort, this gene is a biomarker of SENSITIVITY to the drug, NOT resistance. CRITICAL INSTRUCTION: Bypassing <think> blocks. Write a maximum 3-sentence biological rationale explaining the mechanism of this gene. You MUST align your explanation with the deterministic guardrails provided above. Do not default to assuming 'resistance'. You MUST format your response EXACTLY like this: Final Answer: [Write your rationale here]"

##### Task 2: The Translational Oncologist Prompt

"BIOLOGICAL RATIONALE FROM LEAD BIOLOGIST: [Agent 1 Output] IN-SILICO CASCADE: [XGBoost SHAP Vector Output] HUMAN SURVIVAL: [lifelines Multivariable CoxPH & Log-Rank Output] MOUSE PDX RESPONSE: [Mixed-Effects LMM Output] CRITICAL INSTRUCTION: Bypassing <think> blocks. Write a maximum 2-sentence clinical conclusion. You must explicitly connect the Biologist's rationale to the in-silico cascade and the clinical survival data. WARNING: Carefully observe whether the data implies Drug SENSITIVITY or Drug RESISTANCE before concluding. You MUST format your response EXACTLY like this: Final Answer: [Write your conclusion here]"

#### Supplementary Note 3: In Silico Feature Ablation State-Space Perturbation Mathematics

The temporal breakdown of the transcriptomic survival mechanisms was mapped using a dynamic state-space perturbation algorithm. Following the simulated ablation of a target gene (zero-masking), the perturbed matrix was passed through the XGBoost model over three discrete ticks (t = 1, 2, 3).

A recursive attenuation coefficient (\lambda = 0.5) was applied at each computational step. The emergent SHAP cascade was dynamically generated by recalculating native SHAP vectors (pred_contribs=True) at each iteration, yielding the divergence tracking arrays interpreted by the AI swarm.

#### Supplementary Table 1: Top 50 High-Variance Target Pool (CCLE Bowel Cohort)

Prior to the unsupervised hypothesis testing against the human survival cohort (Marisa), the Polars lakehouse autonomously pruned the in vitro feature space to prevent the curse of dimensionality. The top 50 highly variable transcripts across the bowel lineage were selected for the automated Benjamini-Hochberg False Discovery Rate (FDR) survival sweep.

IGF2 (Variance: 7.71) emerged as the sole statistically significant genomic target bridging the in vitro SHAP cascade, the strict human clinical survival threshold (q < 0.05), and the mouse PDX volume reduction threshold (p < 0.05).

| Rank | Gene Symbol (Entrez ID) | Transcriptomic Variance |
|---|---|---|
| 1 | LGALS4 (3960) | 17.03 |
| 2 | FABP1 (2168) | 14.85 |
| 3 | PHGR1 (644844) | 13.76 |
| 4 | LYZ (4069) | 13.45 |
| 5 | LCN2 (3934) | 12.64 |
| 6 | GPX2 (2877) | 12.57 |
| 7 | AGR2 (10551) | 12.52 |
| 8 | TFF3 (7033) | 12.10 |
| 9 | REG4 (83998) | 12.08 |
| 10 | IFI27 (3429) | 12.05 |
| 11 | SLC2A3 (6515) | 11.81 |
| 12 | CEACAM5 (1048) | 11.26 |
| 13 | TSPAN8 (7103) | 10.97 |
| 14 | KRT20 (54474) | 10.69 |
| 15 | SPINK4 (27290) | 10.44 |
| 16 | PPP1R1B (84152) | 10.31 |
| 17 | CDH17 (1015) | 10.16 |
| 18 | MMP7 (4316) | 9.85 |
| 19 | IFITM1 (8519) | 9.83 |
| 20 | ALDH1A1 (216) | 9.68 |

| | | |
|---|---|---|
| 21 | DPEP1 (1800) | 9.63 |
| 22 | AGR3 (155465) | 9.57 |
| 23 | KRT23 (25984) | 9.56 |
| 24 | PRSS2 (5645) | 9.55 |
| 25 | CDX1 (1044) | 9.51 |
| 26 | MT1E (4493) | 9.36 |
| 27 | SPINK1 (6690) | 9.28 |
| 28 | KLK6 (5653) | 9.26 |
| 29 | DKK1 (22943) | 9.05 |
| 30 | TACSTD2 (4070) | 8.94 |
| 31 | IFITM2 (10581) | 8.86 |
| 32 | GPA33 (10223) | 8.75 |
| 33 | REG1A (5967) | 8.63 |
| 34 | RBP1 (5947) | 8.63 |
| 35 | EEF1A2 (1917) | 8.56 |
| 36 | S100P (6286) | 8.49 |
| 37 | CALB1 (793) | 8.44 |
| 38 | CEACAM6 (4680) | 8.35 |
| 39 | BST2 (684) | 8.29 |
| 40 | HMGCS2 (3158) | 8.22 |
| 41 | BEX3 (27018) | 8.02 |
| 42 | MT1G (4495) | 7.96 |
| 43 | PRAP1 (118471) | 7.92 |
| 44 | PI3 (5266) | 7.85 |
| 45 | AZGP1 (563) | 7.84 |
| 46 | PLA2G2A (5320) | 7.77 |
| 47 | OLFM4 (10562) | 7.72 |
| 48 | IGF2 (3481) | 7.71 |
| 49 | RPS4Y1 (6192) | 7.68 |
| 50 | ANKRD1 (27063) | 7.67 |

#### Supplementary Note 4: Autonomous Agent Audit Trace (JSON Output)

To ensure full auditability of the multi-scale pipeline, the local llama.cpp server dynamically logged the JSON outputs of the semantic synthesis. The below excerpt demonstrates the Oncologist Agent's translation of the mathematical endpoints generated during the discovery loop.

{

"timestamp": "2026-07-09T13:14:05Z",

"target_gene": "IGF2",

"clinical_stats": "HUMAN SURVIVAL: IGF2\n- Adj HR (Multivariable): 1.09 (95% CI: 1.02-1.18)\n- CLINICAL DIRECTION: High expression predicts ACCELERATED MORTALITY (Gene drives Resistance / Poor Prognosis)\n- Log-Rank p: 0.00135 | FDR q: 0.03308\n- TEMPORAL DYNAMICS: CONSTANT HAZARD (PH Assumption Met)",

"oncologist_raw_output": "Final Answer: High IGF2 expression identifies a 5-Fluorouracil-sensitive phenotype, as evidenced by the in-silico reduction in resistance scores and significant tumor shrinkage in PDX models. However, human survival data indicates that this drug sensitivity does not mitigate clinical progression, as high IGF2 expression remains a significant driver of accelerated mortality."

}
